# Molecular typing and macrolide resistance analyses of *Treponema pallidum* in heterosexuals and men who have sex with men communities in Japan, 2017

**DOI:** 10.1101/375485

**Authors:** Mizue Kanai, Yuzo Arima, Shingo Nishiki, Ken Shimuta, Ichiro Itoda, Tamano Matsui, Kazunori Oishi, Makoto Ohnishi, Shu-ichi Nakayama

## Abstract

In recent years, syphilis notifications have increased dramatically in Japan. We performed molecular typing and analyzed macrolide resistance of *Treponema pallidum* samples collected from four clinics and a hospital in Tokyo and Osaka prefectures in 2017.

Macrolide resistant strain type 14d/f was found significantly more in heterosexual syphilis cases compared with those strains identified in men who have sex with men (MSM) syphilis cases. The proportion of 14d/f among heterosexuals was 79% (31/39) compared to 37% (7/19) among MSM [OR, 6.6 (95% CI, 1.7 to 26.7); P=0.002]. 83% (50/60) of the strains were identified as macrolide resistant with an A2058G mutation at the 23S rRNA gene. 90% (35/39) of the heterosexual strains were macrolide resistant, relative to 58% (11/19) of MSM strains; the odds of having the resistant mutation was considerably higher in heterosexuals compared with MSM [OR, 6.4 (95% CI, 1.3 to 33.5); P=0.02]. Heterosexual females and males showed similar distributions, and the results remained the same when restricted to men.

The strain type distribution and frequency of macrolide resistance differed substantially between heterosexual and MSM syphilis samples, suggesting distinct epidemiologic profiles for the two communities and insight into syphilis transmission dynamics in Japan.

## Introduction

With the global re-emergence of syphilis, the number of reports of syphilis has also been rapidly increasing in Japan. Importantly, as with some other countries (1), the recent increase in syphilis case reports in Japan since 2013 has mostly been associated with heterosexual men and women (i.e. men who have sex with women (MSW) and women who have sex with men (WSM)), rather than through transmission amongst men who have sex with men (MSM) (2). This has increased the level of public health concern, given the risk of congenital syphilis (2). Large numbers of syphilis cases have been reported from urban areas such as Tokyo and Osaka prefectures. Molecular typing and macrolide resistance of *Treponema pallidum (T. pallidum)* circulating in Japan have not been frequently or comprehensively reported; importantly, descriptions of the relative distributions of such data according to subpopulations (i.e., heterosexual men and women and MSM) remain unknown. Such information may provide useful insight to the epidemiology of syphilis in Japan, including the current state of macrolide resistance and the dynamics of syphilis transmission. Therefore, we conducted this study to describe the relative frequencies and distributions of the *T. pallidum* strains circulating in Japan, taking into consideration the key subpopulations affected by the current syphilis outbreak.

## Materials and Methods

Specimen samples were collected from four clinics and a hospital specializing in sexually transmitted infections or infectious diseases located in Tokyo and Osaka prefectures in Japan, between January and December 2017. While there was no defined sampling scheme, samples collected were from patients clinically suspected as syphilis by the clinicians. One sample was collected per patient, and these samples were transported to the National Institute of Infectious Diseases laboratory in Tokyo. The majority of samples were swab samples from genital, anal, or oral lesions, and the rest were from cerebrospinal fluid, blood, or surgical samples. Information regarding the case’s gender and self-reported transmission route (i.e., gender of the sex partner(s) suspected as the source of infection) were also collected. For the detection of *T. pallidum* DNA, PCR amplification of *polA* and *tpp47* genes was performed as described previously (3, 4). We considered the sample to be *T. pallidum* DNA positive when PCR amplification was positive for at least one of these two treponemal loci. For *T. pallidum* DNA-positive samples, DNA was extracted from swab samples or isolates using the DNeasy Blood & Tissue Kit (QIAGEN Inc. Valencia, CA.) according to the manufacturer’s protocol, and further evaluations for the enhanced molecular typing and macrolide resistance were conducted. Molecular typing was determined based on the combination of the numbers of 60 bp repeats of the *arp* gene, the restriction fragment length polymorphism (RFLP) patterns of the *tpr* gene, and the sequencing type of the *tp0548* gene. Macrolide resistance was determined by the presence of a mutation in one or both of the 23S rRNA genes (A2058G or A2059G) of *T. pallidum.*

These analyses were performed as described previously (5, 6), except for the modification in the PCR cycling conditions (i.e. annealing/extension steps combined at the single step at 68°C, which used the PCR enzyme, Takara Mighty Amp DNA polymerase (Takara Bio, Inc. Kusatsu, Shiga, Japan)).

Statistical analysis were performed using OpenEpi Version 3.01 (7). While mostly descriptive, a limited number of statistical significance tests were performed to compare the distributions between the MSM and heterosexual samples, based on the two-tailed

Fisher’s exact test; p values less than 0.05 were considered to be statistically significant. To indicate the magnitude of the association, odds ratios (OR) and their associated exact 95% confidence intervals (CI) were also estimated.

This study was approved by the Institutional Review Board of the National Institute of Infectious Diseases (approval number 508 and 705).

## Results

During the study period, a total of 156 samples were collected, and 82 samples (53%) were positive for *T. pallidum* DNA. Positivity varied by transmission route, with 25/53 (47%) MSM samples, 36/56 (64%) heterosexual male samples, and 12/35 (34%) heterosexual female samples testing positive. Among the remaining 12 samples with unknown route of transmission, nine (75%) were positive. Excluding these nine cases, there was a total of 73 samples included in the analysis.

Fully typed profiles were obtained for 58/73 (79%) samples. Four *arp* subtypes (7, 10, 12, 14), ten *tpr* subtypes (a, b, d, e, f, j, k, l, o, p), and six *tp0548* subtypes [a, c, d, f, g, and f- variant 1 (a new type; 1 SNP of type f, G170 is replaced by T, within *tp0548,* deposited to Genbank DDBJ, Accession number LC380837)] were identified. A total of 16 strain types were identified, with 14d/f representing the most common strain type accounting for 38/58 samples (66%), followed by 14d/g (3/58 samples, 5%), 14j/f, 14l/f, and 14p/f (two samples each, 3% each), 7o/c, 10b/a, 10o/c, 12a/c, 14a/f’, 14b/c, 14b/d, 14d/c, 14e/f, 14f/f, and 14k/f (one sample each, 2% each) (Simpson’s diversity index, 0.56).

Macrolide resistance was successfully determined in 60/73 (82%) samples. The majority of these (50/60; 83%) were identified as being macrolide resistant with the A2058G mutation at the 23S rRNA gene; the remaining ten (17%) samples were identified as wild type and susceptible to macrolides. No A2059G mutation at the 23S rRNA gene was identified.

Of 58 fully typed samples, 39 (67%) were from heterosexuals (30 samples from male, nine samples from female), and 19 samples (33%) were from MSM. Six strain types were identified in heterosexuals (five in male, three in female), with 14d/f representing the most common strain type accounting for 31/39 samples [79%; 24/30 (80%) samples in male, and 7/9 (78%) samples in female, Simpson’s diversity index, 0.36]. (Figure 1-A) In contrast, 11 strain types were identified in MSM, with 14d/f being less common [7/19 samples; (37%)], with proportionately more non-14d/f strain types, including 14d/g (3/19 samples; 16%) or other wide spectrum strain types (9/19 samples; 47%) being found in MSMs (Simpson’s diversity index, 0.81). (Figure 1-A) Strain type 14d/f was significantly associated with heterosexual status, and the odds of having this strain type was six times greater in heterosexuals relative to MSM [OR=6.6 (95% CI, 1.7 to 26.7); P=0.002]. The association was the same when the heterosexual group was restricted to men (OR=6.9; 95%CI=1.6-30.6; P=0.006).

**Fig. 1.**
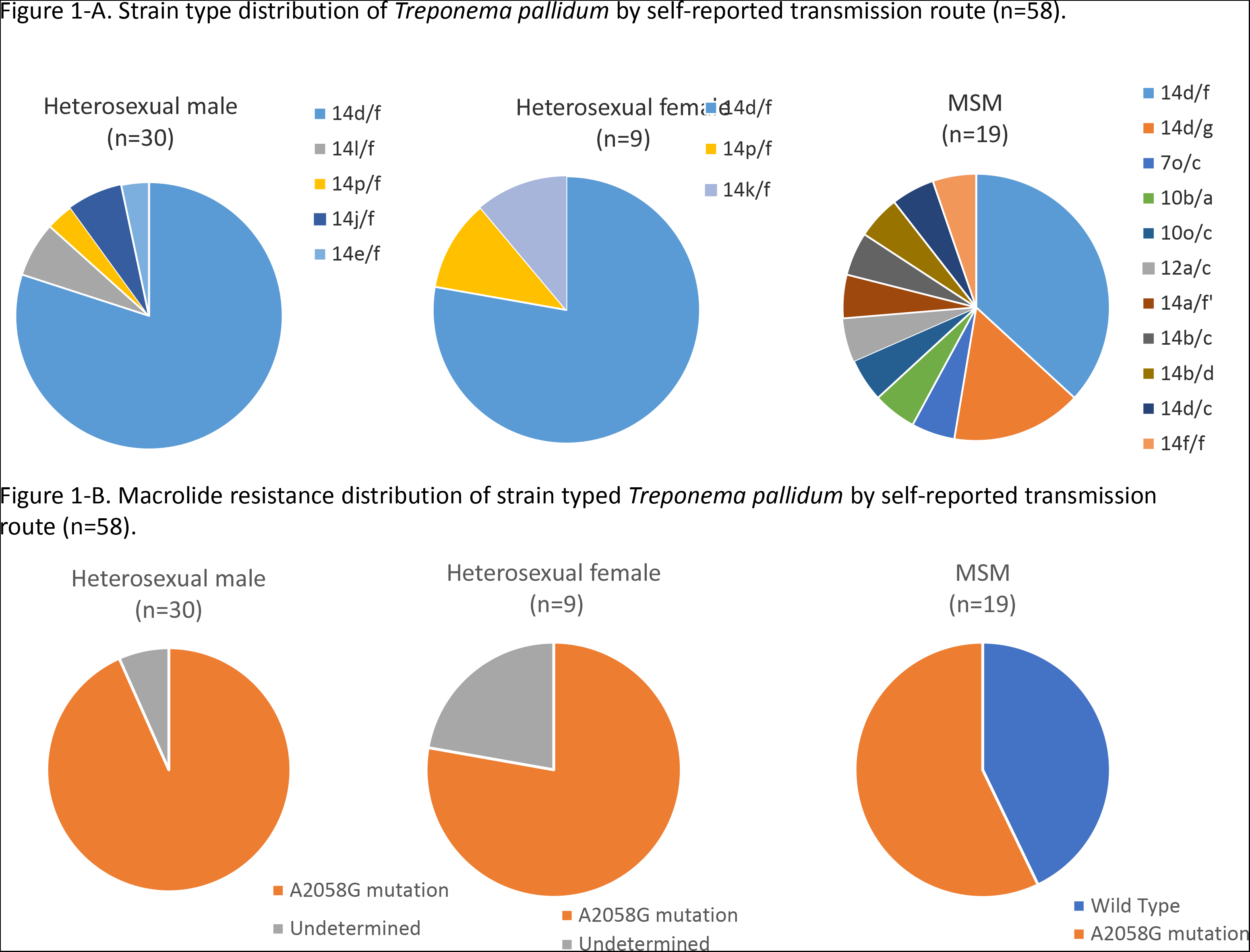
Distributions of molecular types (A) and macrolide resistance (B) among fully typed strains of ***Treponema pallidum*** by self-reported transmission route. A total of 58 fully strain typed samples of *Treponema pallidum* and their macrolide resistance were analyzed by self-reported transmission route (heterosexual male, heterosexual female, and men who have sex with men (MSM)). The strains that 23S rRNA gene could not be determined are indicated as ‘undetermined’ in Figure 1-B.

Almost all fully typed samples in heterosexuals (35/39 samples, 90%) were identified as being macrolide resistant [28/30 samples (93%) in heterosexual males, and 7/9 samples (78%) in female], compared with that in MSM (11/19 samples; 58%). Macrolide resistance was significantly associated with heterosexual status, with a six-fold odds in the mutation being present in heterosexuals relative to MSM [OR=6.4 (95% CI, 1.3 to 33.5); P=0.02]; the association remained unchanged when the heterosexual group was restricted to men (OR=10.2; 95%CI=1.6-107.2; P=0.009). Among MSM, the remaining eight samples (42%) were identified as wild type and susceptible to macrolides. (Figure 1-B).

We conducted further analysis regarding the distribution of macrolide resistance and strain type of *T. pallidum,* for each transmission route group. All of the strain type 14d/f samples in heterosexuals, except for one, were identified as being macrolide resistant (30/31 samples; 97%). Other macrolide resistant strain types in this group included 14e/f, 14j/f, 14l/f, and 14p/f. On the other hand, among MSM, only 4/7 (57%) samples of strain type 14d/f were identified as being macrolide resistant. Among strain type 14d/f samples (n=38), macrolide resistance was significantly associated with heterosexual status [OR, 22.5 (95% CI, 1.3 to 1218); P=0.03]. Overall, among 39 heterosexual samples, 30 (77%) were strain type 14d/f with macrolide resistance, whereas only 4/19 (21%) MSM samples were of this type. Furthermore, all strain type 14d/g strains (3/3 samples) in MSM were identified as being macrolide resistant. Other macrolide resistant strain types in MSM included 7o/c, 10o/c, 12a/c, and 14b/d.

## Discussion

This study reports the first strain typing and macrolide resistance data for *T. pallidum* and its diversity in the heterosexual and the MSM populations in Japan, based on samples from 2017 when the number of syphilis reports continued to increase dramatically among heterosexuals.

Our results revealed that strain type 14d/f with macrolide resistance was the dominant *T. pallidum* strain in Japan, which was similar to previous reports from other countries, such as China (8, 9). The second most common strain type 14d/g, which was seen only in MSM (and the second most common type found in MSM samples), was also previously reported as the dominant strain type in the United States, Australia, and France, where syphilis cases are mainly reported through MSM transmission (6, 10, 11); in Australia, a high proportion of macrolide resistance in strain type 14d/g was also reported (10).

Notably, our results indicated large differences in the distribution of the strain type and macrolide resistance amongst *T. pallidum* in heterosexuals and MSM samples. Heterosexual samples were strongly associated with strain type 14d/f and macrolide resistance, with the co-presence of both being high in this group. In fact, the overall predominance of strain type 14d/f with macrolide resistance was driven by heterosexual samples, with less than a quarter of MSM samples having this molecular characteristic. In addition, MSM lacked such a predominant stain type, displaying a diverse spectrum of various strain types. Restricting to comparisons in males (i.e., heterosexual men vs. MSM), the magnitude of association between heterosexuals and strain type 14d/f and macrolide resistance remained the same. These results suggest that recent *T. pallidum* strains circulating in the heterosexual community in Japan appear to be more homogenous than those circulating in the MSM community, and non-14d/f, non-macrolide resistant strains may continue to circulate in MSM communities. While the number of samples are relatively small, the homogeneity observed in samples from heterosexual men and women during 2017—when reports of MSW and WSM syphilis cases continued to increase considerably—is noteworthy. Although sexual networks can overlap between heterosexual and MSM communities, the differential molecular profile between these two groups indicate that such bridging may be limited, and there may be localized circulation of *T. pallidum* within the heterosexual community that is sustaining the recent increase of syphilis case reports in Japan. Taken together with the strain information from overseas, our results suggest that the transmission dynamics, including sexual networks between Japan and other countries, may differ between the heterosexual and MSM communities in Japan.

There are some limitations in this study regarding the geographical origin of the samples obtained. As the samples were collected from Tokyo and Osaka prefectures, our results may not be generalizable to all the strain characteristics of *T. pallidum* across Japan. However, as these two prefectures have the highest syphilis burden in Japan (for both absolute number of notifications and notification rate per population) (2), they warrant particular focus; in addition, with the majority of MSM cases being reported from these urban areas, a more controlled comparison of the strains can be made between the MSM and heterosexual samples, as the geographic location from where the cases arose from are the same. Another limitation is with regarding the self-reported nature of the transmission route (i.e. heterosexual vs. MSM). Given the societal context in Japan, misclassifications of MSM as heterosexual men for social desirability reasons is possible (i.e., MSM misreporting themselves as heterosexual), but such bias would lead to an underestimate in the observed associations; in fact, the large contrast in the molecular profile observed between the two groups provide reassuring evidence regarding the validity of the self-reported information.

In conclusion, this study found important differences in the strain type and macrolide resistance of *T. pallidum* recently circulating in the heterosexual and MSM communities in Japan. Together with epidemiologic information, these molecular findings help to provide a better understanding of the transmission dynamics of the syphilis epidemic—not only within Japan but also within the broader global context—which can in turn help inform public health response for syphilis control.

## Acknowledgements

The authors are grateful to Igen Hongo, Masayuki Sawamura, Takashi Hamada, and Hiroshi Kameoka who generously and continuously provided us with specimens from clinically suspected syphilis cases. The authors also thank Kimiko Matsumoto for her technical support in the detection and typing of *T. pallidum,* including PCR procedures. This study was supported by a grant from the Ministry of Health, Labour and Welfare of Japan (H-28-Shinkou-Gyousei-Ippan-008).

## References

1. Kamb ML, Taylor MM, Ishikawa N. 2018. Rapid increases in syphilis in reproductive-aged women in Japan:A warning for other countries? Sex. Transm. Dis. 45:144–146.

2. Takahashi T, Arima Y, Yamagishi T, Nishiki S, Kanai M, Ishikane M, Matsui T, Sunagawa T, Ohnishi M, Oishi K. 2018. Rapid increase in reports of syphilis associated with Men Who Have Sex with Women and Women Who Have Sex with Men, Japan, 2012 to 2016. Sex. Transm. Dis. 45:139–143.

3. Liu H, Rodes B, Chen C Y, Steiner B. 2001. New tests for syphilis: rational design of a PCR method for detection of *Treponema pallidum* in clinical specimens using unique regions of the DNA polymerase I gene. J. Clin. Microbiol. 39:1941–1946.

4. Orle KA, Gates CA, Martin DH, Body BA, Weiss JB. 1996. Simultaneous PCR detection of *Haemophilus ducreyi, Treponema pallidum*, and herpes simplex virus types 1 and 2 from genital ulcers. J. Clin. Microbiol. 34:49–54.

5. Marra C, Sahi S, Tantalo L, Godornes C, Reid T, Behets F, Rompalo A, Klausner JD, Yin Y, Mulcahy F, Golden MR, Centurion-Lara A, Lukehart SA. 2010. Enhanced molecular typing of *treponema pallidum*: geographical distribution of strain types and association with neurosyphilis. J. Infect. Dis. 202:1380–1388.

6. Lukehart SA, Godornes C, Molini BJ, Sonnett P, Hopkins S, Mulcahy F, Engelman J, Mitchell SJ, Rompalo AM, Marra CM, Klausner JD. 2004. Macrolide resistance in *Treponema pallidum* in the United States and Ireland. N. Engl. J. Med. 351:154–158.

7. Dean AG, Sullivan KM, Soe MM. OpenEpi: Open Source Epidemiologic Statistics for Public Health, Version. www.OpenEpi.com, updated 2013/04/06, accessed 2018/03/09.

8. Dai T, Li K, Lu H, Gu X, Wang Q, Zhou P. 2012. Molecular typing of *Treponema pallidum:a* 5-year surveillance in Shanghai, China. J. Clin. Microbiol. 50:3674–3677.

9. Chen XS, Yin YP, Wei WH, Wang HC, Peng RR, Zheng HP, Zhang JP, Zhu BY, Liu QZ, Huang SJ. 2013. High prevalence of azithromycin resistance to *Treponema pallidum* in geographically different areas in China. Clin. Microbiol. Infect. 19:975–979.

10. Read P, Tagg KA, Jeoffreys N, Guy RJ, Gilbert GL, Donovan B. 2016. *Treponema pallidum* strain types and association with macrolide resistance in Sydney, Australia:new *tp0548* gene types identified. J. Clin. Microbiol. 2016. 54:2172–2174.

11. Grange PA, Allix-Beguec C, Chanal J, Benhaddou N, Gerhardt P, Morini JP, Deleuze J, Lassau F, Janier M, Dupin N. 2013. Molecular subtyping of *Treponema pallidum* in Paris, France. Sex. Transm. Dis. 40:641–644.

